# Endogenous tagging reveals differential regulation of Ca^2+^ channels at single AZs during presynaptic homeostatic potentiation and depression

**DOI:** 10.1101/240051

**Authors:** Scott J. Gratz, Pragya Goel, Joseph J. Bruckner, Roberto X. Hernandez, Karam Khateeb, Gregory T. Macleod, Dion Dickman, Kate M. O’Connor-Giles

## Abstract

Neurons communicate through Ca^2+^-dependent neurotransmitter release at presynaptic active zones (AZs). Neurotransmitter release properties play a key role in defining information flow in circuits and are tuned during multiple forms of plasticity. Despite their central role in determining neurotransmitter release properties, little is known about how Ca^2+^ channel levels are modulated to calibrate synaptic function. We used CRISPR to tag the *Drosophila* Ca_V_2 Ca2+ channel Cacophony (Cac) and investigated the regulation of endogenous Ca^2+^ channels during homeostatic plasticity in males in which all endogenous Cac channels are tagged. We found that heterogeneously distributed Cac is highly predictive of neurotransmitter release probability at individual AZs and differentially regulated during opposing forms of presynaptic homeostatic plasticity. Specifically, Cac levels at AZ are increased during chronic and acute presynaptic homeostatic potentiation (PHP), and live imaging during acute expression of PHP reveals proportional Ca^2+^ channel accumulation across heterogeneous AZs. In contrast, endogenous Cac levels do not change during presynaptic homeostatic depression (PHD), implying that the reported reduction in Ca^2+^ influx during PHD is achieved through functional adaptions to pre-existing Ca^2+^ channels. Thus, distinct mechanisms bi-directionally modulate presynaptic Ca^2+^ levels to maintain stable synaptic strength in response to diverse challenges, with Ca^2+^ channel abundance providing a rapidly tunable substrate for potentiating neurotransmitter release over both acute and chronic timescales.

## Introduction

Dynamic changes in the properties controlling neurotransmitter release at individual synaptic connections modulate information flow in neural circuits. Neurotransmitter release occurs at specialized domains called active zones (AZs) where synaptic vesicles fuse to presynaptic membranes and release their contents following an action potential-driven influx of Ca^2+^. AZ release properties are determined locally and can vary considerably within and between neuronal subtypes (Hatt and Smith, 1976; Atwood and Karunanithi, 2002; Branco and Staras, 2009; Sudhof, 2012). However, how this remarkable diversity in AZ release properties is established, maintained, and dynamically adjusted during various forms of plasticity is poorly understood.

Synaptic vesicle release is a stochastic process that is subject to dynamic modulation over both acute and chronic timescales. The probability that an action potential will elicit fusion of a synaptic vesicle at a particular AZ (synaptic probability of release, P_r_) is a defining property of neurotransmitter release that can be tuned during plasticity. Two key parameters have a large influence on P_r_: the number of synaptic vesicles available for release and their individual probability of release (P_v_). P_v_ is heavily influenced by Ca^2+^ influx through presynaptic Ca^2+^ channels, which in turn is driven by the number, organization, and intrinsic properties of Ca^2+^ channels at AZs (Ariel and Ryan, 2012; Ehmann et al., 2015; Dittrich et al., 2018).

The *Drosophila* neuromuscular junction (NMJ) provides a powerful model for investigating the establishment and modulation of release properties at individual AZs *in vivo*. At fly NMJs, single *Drosophila* glutamatergic motorneurons form hundreds of synapses with their postsynaptic muscle targets. These synapses have many presynaptic parallels to excitatory synapses in the mammalian CNS, including molecular composition, ultrastructural organization of AZs, and neurotransmitter release properties (Ackermann et al., 2015; Harris and Littleton, 2015; Zhan et al., 2016; Van Vactor and Sigrist, 2017). Further, functional imaging studies with genetically encoded Ca^2+^ indicators have facilitated the investigation of neurotransmission at single synapses of *Drosophila* motorneurons, revealing a remarkable degree of heterogeneity in release properties between individual AZs (Guerrero et al., 2005; Peled and Isacoff, 2011; Melom et al., 2013; Peled et al., 2014; Akbergenova et al., 2017). Finally, conserved forms of homeostatic plasticity are expressed at the *Drosophila* NMJ. In a process known as presynaptic homeostatic potentiation (PHP), genetic or pharmacological disruption of postsynaptic glutamate receptors induces a compensatory increase in neurotransmitter release that maintains overall synaptic strength (Frank, 2014; Davis and Muller, 2015). An opposing form of homeostatic plasticity, called presynaptic homeostatic depression (PHD) is observed in response to excess glutamate released from individual synaptic vesicles. In this paradigm, a homeostatic decrease in the number of vesicles released stabilizes synaptic strength (Daniels et al., 2004). While the two forms of homeostatic plasticity rely on inverse modulations to Ca^2+^ influx through presynaptic Ca^2+^ channels (Muller and Davis, 2012; Gavino et al., 2015), the mechanisms that achieve these adaptations are unclear.

Here, we engineered the endogenous locus of *Drosophila cacophony (cac)* to incorporate a fluorescent tag in all 15 known isoforms of Cac. Cac is the pore-forming subunit of the sole Ca^2+^ channel required for triggering synchronous neurotransmitter release in *Drosophila* (Smith et al., 1996; Kawasaki et al., 2000; Macleod et al., 2006; Peng and Wu, 2007). The generation of endogenously tagged Cac allowed us to probe the relationship between Cac levels, P_r_, and homeostatic plasticity. We found that the abundance of endogenous Cac is modulated during PHP, but does not change during PHD, indicating that distinct mechanisms are employed to bidirectionally tune Ca^2+^ influx during these opposing forms of homeostatic plasticity. Live imaging of Cac at identified synapses before and after acute pharmacological induction of PHP revealed the proportional accumulation of Ca^2+^ channels across AZs, a mechanism that maintains the heterogeneous distribution of Ca^2+^ channels. We propose that differential regulation of Ca^2+^ channels at single AZs confers reliable neurotransmission and a broad capacity for tuning neurotransmitter release probability to maintain neural communication.

## Materials and methods

### *Drosophila* genetics and genome engineering

The following fly lines are available at the Bloomington *Drosophila* Stock Center (BDSC): *w*^*1118*^ (BDSC #5905), *vasa-Cas9* (BDSC #51324), and piggyBac transposase (BDSC #8285) (Horn et al., 2003; Parks et al., 2004; Gratz et al., 2014). The null allele *GluRIIA*^*SP16*^ (Petersen et al., 1997) and *UAS-vGlut* (Daniels et al., 2004) were generously provided by Aaron DiAntonio. PS-GCaMP comprises an MHC promoter for muscle expression, an N-terminal myristoylation sequence for membrane targeting, GCaMP5, and the Shaker PDZ domain for targeting to the postsynaptic density, and was incorporated into the attP2 landing site (BDSC-25710).

Endogenously tagged Cacophony was generated using a scarless CRISPR/piggyBac-based approach (flyCRISPR.molbio.wisc.edu) (Bruckner et al., 2017). Briefly, sequences coding for sfGFP or TagRFP-T flanked by flexible linkers and a visible marker flanked by piggyBac inverted terminal repeat sequences were inserted immediately downstream of the transcriptional start site of the endogenous *Cacophony* locus (basepairs 11,980,057 through 11,980,055, *D. melanogaster* genome release 6). This site is in an exon common to all isoforms of *Cac.* piggyBac transposase was subsequently used for footprint-free removal of the marker, followed by molecular confirmation of precise tag incorporation.

### Presynaptic Ca^2+^ imaging and analysis

Ca^2+^ influx at motor terminals was measured by forward filling of dextran-conjugated indicators (Macleod, 2012). Male third instar larvae were dissected in chilled Schneider’s insect medium (Sigma). Severed segmental nerves were drawn into a filling pipette, and a 16:1 mixture of the Ca^2+^ indicator rhod dextran (R34676; Thermo Fisher Scientific) and a Ca^2+^-insensitive Alexa Fluor 647 dextran (AF647 dextran; D22914; Thermo Fisher Scientific) was applied to the cut nerve end for 15-45 minutes. Preparations were incubated in the dark for at least 3 hours and rinsed every 30 minutes with fresh Schneider’s insect medium. Thirty minutes before imaging, the Schneider’s insect medium was replaced with hemolymph-like solution (HL6) containing 2.0 mM Ca^2+^ and supplemented with 7 mM L-glutamic acid (Macleod et al., 2002). Analysis was performed on type-Ib and type-Is motor neuron terminals innervating muscle 6 in hemisegment A3 or A4. Live imaging was performed on a Nikon Eclipse FN1 microscope with a 100x 1.1NA water immersion objective (Nikon) using an iXon3 888 EMCCD camera (Andor) operating at a frame rate of 114 Hz. The terminals were alternately excited at 550 ± 7nm and 640 ± 15nm, and emission was alternately collected at 605 ± 26nm and 705 ± 36nm, respectively. The cut end of the segmental nerve was stimulated using a 0.4-ms pulse 10 times at 1 Hz followed by a stimulus train at 20 Hz for one second. Images were background subtracted and analyzed by generating fluorescence intensity traces with NIS-Elements Ar (Nikon). For each preparation (N), 2-5 non-terminal boutons of a single NMJ were analyzed. Ca^2+^ levels are measured as the fluorescence ratio of rhod to AF647. Ten single action potentials were averaged into a single trace and used to calculate peak amplitude and the decay time constant (tau). Ca^2+^ imaging data were excluded from further analysis if they were collected from terminals where resting Ca^2+^ levels were assessed to be outliers. The bounds used to assess outliers were calculated as follows: the Median Absolute Deviation (MAD) was calculated for resting Ca^2+^ levels of each group of terminals and the upper boundary was defined as 3x MAD above the median, and the lower boundary was defined as 3x MAD below the median (Leys, 2013). No further criteria for exclusion of Ca^2+^ imaging data were applied, providing the fluorescence transient of the Ca^2+^ indicator recovered to the baseline with a time course of < 250ms (tau).

### Electrophysiology

Male third instar larvae were dissected in modified Ca^2+^-free hemolymph-like saline (HL3, 70 mM NaCl, 5 mM KCl, 10 mM MgCl2, 10 mM NaHCO3, 115 mM sucrose, 5 mM trehalose, 5 mM HEPES, pH 7.2) (Stewart et al., 1994) as described (Kiragasi et al., 2017). Briefly, neuromuscular junction sharp electrode (electrode resistance between 10-30 MΩ) recordings were performed in HL3 saline containing 0.4 mM Ca^2+^ on muscles 6 and 7 of abdominal segments A2 and A3 using a sharp borosilicate electrode (resistance of 15-25 MΩ) filled with 3M KCl. Recordings were performed on an Olympus BX61 WI microscope using a 40x/0.80 water-dipping objective and acquired using an Axoclamp 900A amplifier, Digidata 1440A acquisition system and pClamp 10.5 software (Molecular Devices). Electrophysiological sweeps were digitized at 10 kHz, and filtered at 1 kHz.

Miniature excitatory junctional potentials (mEJPs) were recorded in the absence of any stimulation, and cut motor axons were stimulated to elicit excitatory junctional potentials (EJPs). For each recording, at least 100 mEJPs were analyzed using Mini Analysis (Synaptosoft) to obtain a mean mEJP amplitude value. EJPs were stimulated with an ISO-Flex stimulus isolator (A.M.P.I.), with intensity adjusted for each cell to consistently elicit responses from both type Ib and Is motor neurons innervating the muscle segment. At least 20 consecutive EJPs were recorded for each cell and analyzed in pClamp to obtain mean amplitude. Quantal content was estimated for each recording by calculating the ratio of mean EPSP amplitude to mean mEPSP amplitude and then averaging recordings across all NMJs for a given genotype. Muscle input resistance (R_in_) and resting membrane potential (V_rest_) were monitored during each experiment. Recordings were analyzed only if the V_rest_ was between -60 mV and -80 mV, and if the R_in_ was ≥ 5MΩ.

Pharmacological homeostatic challenge was assessed by incubating semi-intact preparations in 20 µM Philanthotoxin-433 (Santa Cruz, sc-255421, Lot B1417) diluted in HL3 containing 0.4 mM Ca^2+^ for 10 min at room temperature (Frank et al., 2006). Following treatment, the dissection was completed and the prep was rinsed 5 times in recording solution. For analysis of Ca^2+^ channel levels following acute homeostatic plasticity, preparations were fixed in 4% PFA for 30 minutes following PhTx incubation and rinses, and stained as described below.

### Immunostaining

Male third instar larvae were dissected in ice-cold Ca^2+^-free saline and fixed for 30 minutes in 4% paraformaldehyde in PBS or 5 minutes in 100% ice-cold ethanol. Dissected larvae were washed and permeabilized in PBS containing 0.1% Triton-X and blocked for one hour in 5% normal donkey serum or overnight at 4°C in PBS containing 0.1% Triton-X and 1% BSA. Dissected larvae were incubated in primary antibodies overnight at 4°C or 3 hours at room temperature and secondary antibodies for 2-3 hours at room temperature, then mounted in Vectashield (Vector Laboratories) or ProLong Diamond (ThermoFisher). The following antibodies were used at the indicated concentrations: mouse anti-Brp at 1:100 (Nc82; developed by Erich Buchner and obtained from the Developmental Studies Hybridoma Bank), rabbit anti-GFP conjugated to AlexaFluor 488 at 1:500 (#A21311, ThermoFisher), and anti-HRP conjugated to AlexaFluor 647 at 1:200-1:500 (Jackson ImmunoResearch). Species-specific Alexa Fluor 488 and 568 secondary antibodies (Invitrogen, Jackson ImmunoResearch) were used at 1:400-1:500.

### Confocal imaging and analysis

Confocal images were acquired on a Nikon A1R-Si+ with Plan-Apo 60x 1.40 NA and 100x APO 1.4NA oil immersion objectives or a Olympus Fluoview FV1000 with Plan-Apo 60x (1.42 NA) oil immersion objective. Image analyses and brightness and contrast adjustments were performed using the Fiji distribution of ImageJ (Schindelin et al., 2012). For analysis of Cac and Brp intensity, all genotypes were stained together and imaged in the same session with identical microscope settings optimized for detection without saturation of the signal. For consistency, analysis was limited to type Ib synapses of NMJ 6/7 in segments A2 and A3. Maximum intensity projections were used for quantitative image analysis using either NIS Elements software General Analysis toolkit or Fiji as follows. To measure Cac^sfGFP-N^ and Brp intensity at individual AZs, nonsynaptic structures including axons were removed from the images using freehand selection and fill. Z-stacks were flattened using the Maximum Intensity Z-projection function. Channels were separated, background subtracted, and noise was reduced using a light Gaussian filter (0.75-pixel sigma radius). A threshold was applied to the Brp channel to remove irrelevant low intensity pixels and individual puncta were identified and segmented using the Find Maxima tool. The resulting segmented Brp image was used to create a binary mask that was also used to segment the Cac^sfGFP-N^ channel. Intensity data were collected using the Analyze Particles tool. The fluorescent intensity of each puncta is measured as the SUM fluorescence calculated directly or as the product of average intensity and particle area.

### Live imaging

Functional imaging experiments were performed using a Nikon A1R+ scanning confocal system built on a Nikon Eclipse FN1 microscope with a CFI Plan 100XW objective (Nikon). Cac^TagRFP-N^; PS-GCaMP male third instar larvae were dissected in ice-cold Ca^2+^-free HL3 and severed segmental nerves were drawn into a stimulating pipette. Preparations were then maintained in HL3 saline containing 1.5 mM Ca^2+^ and 25 mM Mg^2+^ for imaging. Type-Ib motor neuron terminals innervating muscle 6 in hemisegment A2 or A3 were first imaged using the galvanometer scanner to collect a Cac^TagRFP-N^ Z-stack. GCaMP5 was then imaged continuously in a single focal plane using the resonant scanner with full frame averaging for a final acquisition rate of 15.3 frames per second. Evoked postsynaptic Ca^2+^ transients were stimulated by applying a 0.4-ms pulse every second. The stimulus amplitude was adjusted to reliably recruit the Ib input. 94-100 stimulations were recorded for each experiment. Cac^TagRFP-N^ puncta location and intensity was analyzed in Fiji. Z-stacks were flattened using the Maximum Intensity Z-projection function. To identify individual Cac^TagRFP-N^ puncta, a mask was created using a gaussian filter (sigma radius = 2) and unsharp mask (radius = 3, weight = 0.8). A threshold was then applied to the mask to remove lower intensity pixels between puncta and individual puncta were segmented using the Find Maxima tool. By identifying large puncta with two local intensity maxima, the Find Maxima tool facilitates the identification and segmentation of closely spaced AZs. From each resulting image, a binary mask was then created and used to isolate puncta in the original Z-projection. Each punctum was then interrogated for intensity and XY coordinates using the Analyze Particles tool. GCaMP movies were processed using Nikon Elements software. Motion artifacts during acquisition were corrected by aligning all frames to the first frame of the movie. Baseline fluorescence was subtracted from every frame using a frame created from the average of 10 non-stimulus frames that also lacked spontaneous release events, and noise was reduced using the Nikon Elements Advanced Denoising function. The XY coordinates of postsynaptic Ca^2+^ transients were collected from all stimulation frames manually or automatically. For manual analysis, XY coordinates were obtained using hand selection of individual fluorescence peaks. Only peaks that persisted and decayed over subsequent frames were recorded. For automated analysis, all stimulation frames were thresholded to remove background noise and the Find Maxima tool in Fiji was used to identify event XY coordinates. Each postsynaptic event was then assigned to a Cac^TagRFP-N^ punctum through nearest neighbor analysis using euclidean distance.

Live imaging of Cac^sfGFP-N^ during acute PHP was performed on a Nikon A1R-HD scanning confocal system built on a Nikon Eclipse FN1 microscope with a CFI75 Apochromat 25x 1.1 NA objective (Nikon). Third instar larva were minimally dissected in Ca^2+^-free HL3 and one side of the body wall was carefully pinned down while avoiding any stretching of body wall muscles. Before treatment, Z-stacks of NMJ 6/7 were acquired using the resonant scanner with 4x averaging. Preparations were then exposed to 40 µM Philanthotoxin-433 (Santa Cruz, sc-255421, Lot F2018) or vehicle diluted in HL3 for 10 min. Because segmental nerves were left intact, experiments were performed in Ca^2+^-free HL3 to minimize muscle contractions and movement during imaging, a condition that does not affect PHP (Goel et al., 2017). Immediately following PhTx treatment, NMJS were reimaged using the same imaging parameters. To measure Cac^sfGFP-N^ intensity at individual AZs, Z-stacks were flattened using the Maximum Intensity Z-projection function and background subtracted. A Gaussian filter (0.9-pixel sigma radius) was applied to the after treatment images to aid in AZ recognition and masking. After images were then segmented using freehand selection and fill to remove nonsynaptic structures. A threshold was applied to remove irrelevant low intensity pixels and individual AZ puncta were identified and segmented using the Find Maxima tool. The resulting segmented image was used to create a binary mask that was used to segment the unfiltered before-and after-treatment background subtracted maximum intensity projections. Intensity data were collected using the Analyze Particles tool and post-PhTx intensities were corrected for vehicle-only effects. For consistency, analysis was restricted to type-Ib AZs of NMJ 6/7.

### Experimental Design and Statistical Analysis

Statistical analyses were conducted in GraphPad Prism 7, SigmaStat 3.5, and R. Single comparisons of normally distributed datasets, as determined by the D’Agostino-Pearson omnibus test, were conducted by Student’s t test. Welch’s correction was used in cases of unequal variance. The Mann-Whitney U test was used for single comparisons of non-normally distributed data. For multiple comparisons of normally distributed data, we performed ANOVA followed by Tukey’s test. One-dimensional Pearson correlation coefficients (r) were used to compare intensity levels and release probability. Fisher’s exact test was used to compare proportions. Reported values are mean ± SEM unless otherwise stated. P values, statistical test used, and sample sizes are also reported in each figure legend.

## Results

### Endogenous tagging of the Ca_V_2 Ca2+ channel Cacophony

Cac is the only Ca_V_2-family member in *Drosophila*, and is homologous to mammalian N-and P/Q-type voltage-gated Ca^2+^ channels (Smith et al., 1996; Littleton and Ganetzky, 2000). Neurotransmitter release depends on the synaptic localization of Ca_V_2 channels and mutations that impinge on their function are associated with significant neurophysiological dysfunction at synapses (Smith et al., 1998; Mochida, 2018; Nanou and Catterall, 2018). Investigation of synaptic Ca^2+^ channel levels and localization in *Drosophila* has relied on overexpression of a C-terminally tagged transgene using the GAL4/UAS system (Kawasaki et al., 2004). This approach has enabled the identification of multiple regulators of Ca^2+^ channel localization to AZs (Kittel et al., 2006; Graf et al., 2009; Liu et al., 2011; Graf et al., 2012; Muller et al., 2012; Bruckner et al., 2017). However, several aspects of this transgenic approach pose limitations for investigating the relationship between Ca^2+^ channels and synapse-specific release properties: (1) *cac* is necessarily overexpressed, generally at high levels, under the control of GAL4 instead of endogenous regulatory elements; (2) only one of 15 *cac* isoforms is expressed; (3) the C-terminal position of GFP likely obstructs a conserved PDZ-binding domain that mediates known interactions with synaptic proteins; and (4) expression of the single Cac isoform was shown to rescue viability, but not flight, in null alleles, indicating channel function is not fully restored or properly regulated (Kawasaki et al., 2002; Kawasaki et al., 2004).

To overcome these limitations and establish a reagent for following all endogenous Ca_V_2 channels *in vivo*, we targeted the translational start site of 14 out of 15 *cac* isoforms for incorporation of superfolder-GFP (sfGFP) using CRISPR-based gene editing (Fig. 1A). This insertion site also results in the in-frame translation of sfGFP within the longer N-terminal cytoplasmic domain of the single isoform (isoform N) with an earlier start site, allowing us to follow the full complement of endogenously expressed Cac isoforms (Fig. 1B). *cac*^*sfGFP-N*^ flies are homozygous and hemizygous viable, emerge at expected frequencies, and are capable of effective flight. Cac^sfGFP-N^ localizes to AZs marked by the AZ cytomatrix protein Brp at presynaptic terminals of larval motorneurons with remarkable specificity (Fig. 1C). Endogenous Cac also localizes to the synaptic neuropils of the larval ventral ganglion and adult brain (Fig. 1D,E). In the adult brain, it was immediately apparent that endogenous Cac levels vary between brain regions. Most notably, we consistently observe higher Cac levels in the mushroom body. Therefore, incorporation of a fluorescent tag at the endogenous locus of Cac reports the expected gross localization of Ca^2+^ channels in the *Drosophila* nervous system without obvious defects in viability or behavior.

**Figure 1.**
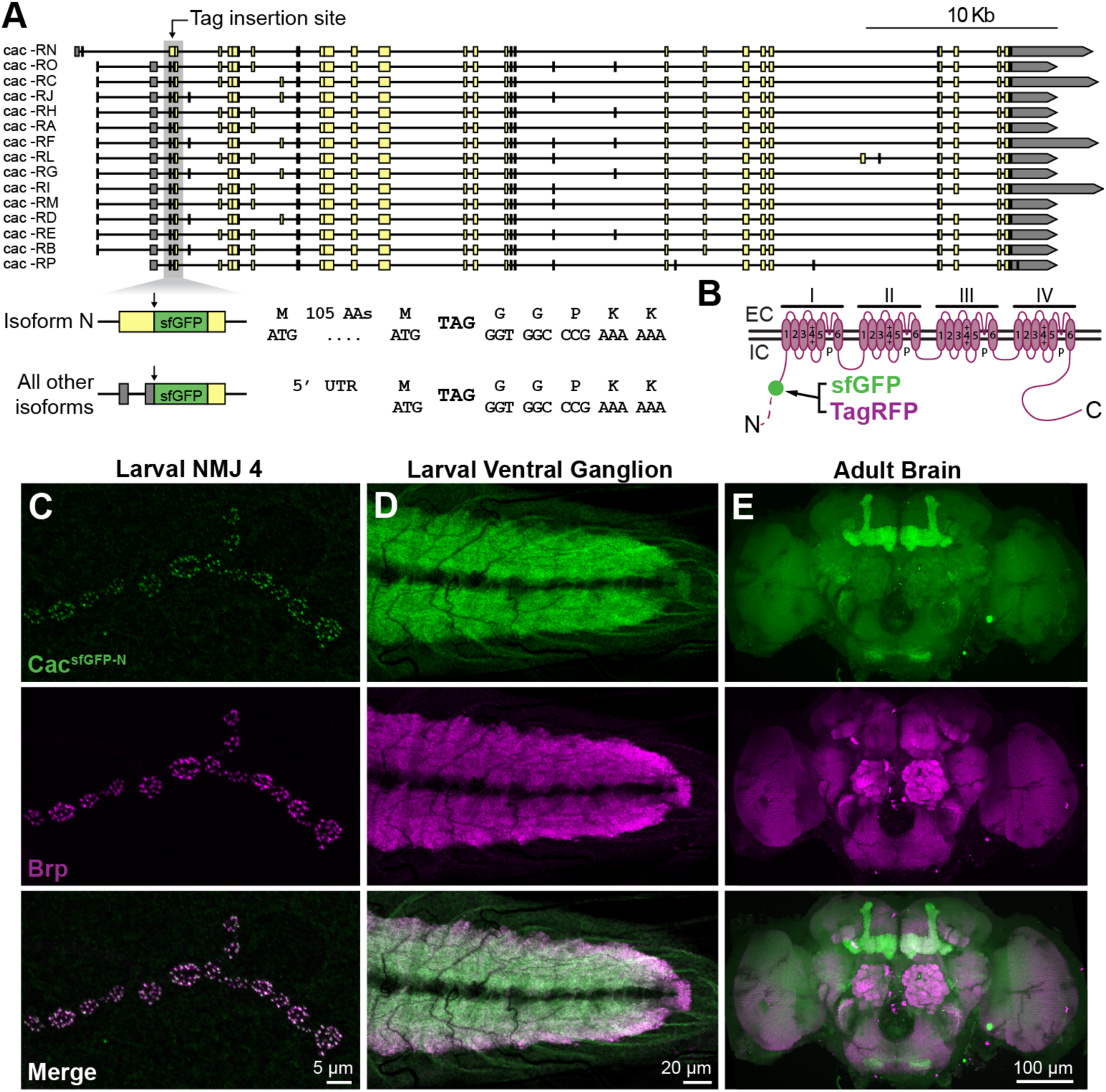
Endogenous tagging of the Ca_V_2 Ca2+ channel Cacophony. **(A)** Map of the *cacophony* locus indicating the genomic region targeted to incorporate sfGFP in all predicted isoforms. sfGFP is inserted between amino acids 107 and 108 of isoform N and immediately following the start codon in all other isoforms. **(B)** Schematic of the endogenously tagged Cacophony protein indicating the site of tag incorporation in the encoded channel (green dot). The longer N-terminus of isoform N is represented by a dotted line. EC – extracellular, IC – intracellular. (**C-E**) Confocal Z-projections of a *cac*^*sfGFP-N*^ larval NMJ (**C**), larval ventral ganglion (**D**), and adult brain (**E**) co-labeled with antibodies against GFP and the active zone marker Brp.

### Normal synaptic function at *Cac*^*sfGFP-N*^ NMJs

To directly evaluate the function of endogenously tagged channels, we first assayed presynaptic Ca^2+^ influx via fluorescence imaging of a Ca^2+^-sensitive dye loaded into axon terminals *in situ* (Macleod, 2012). The use of a rapid-binding chemical dye loaded in fixed proportion to a Ca^2+^-insensitive dye provided ratiometric resolution of single action potential-mediated Ca^2+^ transients and allowed for direct comparisons between terminal types, preparations, and genotypes (Fig. 2A-E) (Lu et al., 2016). The resting level of Ca^2+^ is no different between *cac*^*sfGFP-N*^ and wild-type larvae in either type-Ib (big) or -Is (small) terminals on muscle fiber 6 (Fig. 2C). Similarly, there is no indication of a deficit in single action potential-mediated Ca^2+^ influx in *cac*^*sfGFP-N*^ on the basis of 1 Hz stimuli-evoked fluorescence transients (Fig. 2D). Finally, we measured the amplitude of the Ca^2+^ plateau during trains of action potentials and did not detect a significant change in *cac*^*sfGFP-N*^ relative to wild type (Fig. 2E). Thus, while we cannot rule out the possibility that tag incorporation could have subtle effects on Ca^2+^ channel function, we observe normal Ca^2+^ influx at motor terminals. To further interrogate the impact of endogenous tagging, we investigated synaptic transmission at *cac*^*sfGFP-N*^ NMJs. Consistent with normal Ca^2+^ influx, this analysis revealed no differences between control and *cac*^*sfGFP-N*^ in mEJP frequency, mEJP amplitude, or quantal content (Fig. 2F-K).

In addition to its role in neurotransmission, Cac promotes synapse formation (Rieckhof et al., 2003). We assessed synaptic growth in *cac*^*sfGFP-N*^ animals and found a mild decrease in bouton number at NMJ6/7 (Fig. 2-1A-C). At NMJ4, there is no difference in bouton number between *cac*^*sfGFP-N*^ and wild type (Fig. 2-1A,B,D). Similarly, AZ number per bouton is unaltered at *cac*^*sfGFP-N*^ NMJs (Fig. 2-1E). These analyses, together with the findings described below, indicate that the incorporation of a fluorescent tag into the endogenous protein does not substantively disrupt Cac function *in vivo*.

**Figure 2.**
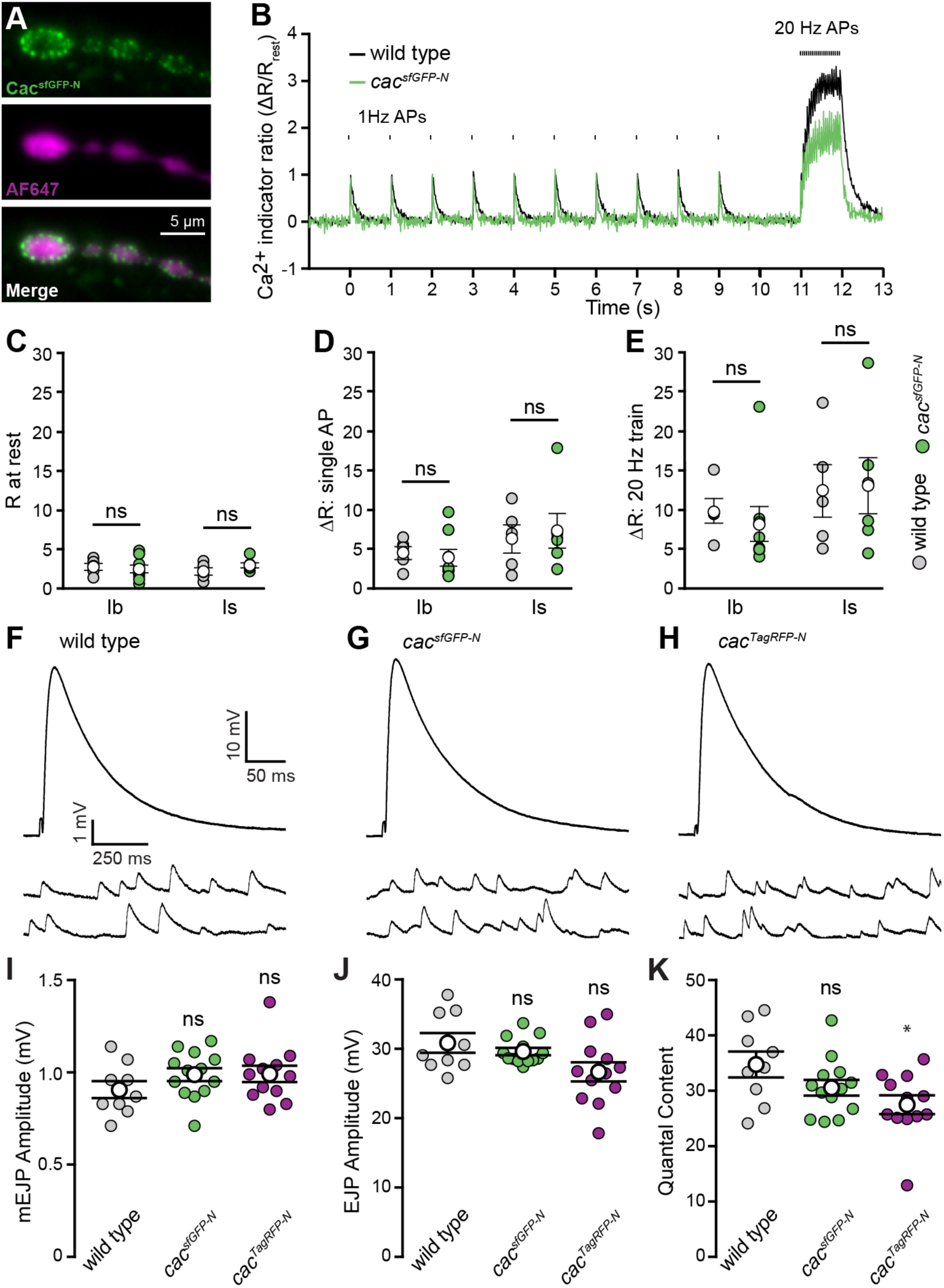
Endogenous tagging of Cacophony does not perturb presynaptic Ca^2+^ influx or synaptic function. **(A)** Live fluorescence images of type-Ib boutons from a *cac*^*sfGFP-N*^ motor terminal showing Cac^sfGFP-N^ (top) and AF647-dextran (center), which was co-loaded with rhod-dextran (not shown). **(B)** Ratiometric fluorescence changes (rhod-dextran relative to AF647-dextran) in presynaptic motorneuron terminals during stimulation in wild type and *cac*^*sfGFP-N*^. An action potential was initiated every second, for a period of 10 seconds, followed by a 1-sec, 20-Hz train of action potentials. **(C)** Plot of average Ca^2+^ levels (R) in terminals prior to nerve stimulation for type-Ib and -Is terminals in wild-type and *cac*^*sfGFP-N*^ boutons (wild-type Ib, 2.74 ± 0.44, one NMJ from each of 5 animals (N=5); *cac*^*sfGFP-N*^ Ib, 2.47 ± 0.49, N=8, p=0.69; wild-type Is, 2.16 ± 0.47, N=5; *cac*^*sfGFP-N*^ Is, 2.94 ± 0.32, N=6, p=0.21; Student’s t test). **(D)** Plot of the average amplitude of single action potential-mediated Ca^2+^ transients (ΔR) (wild-type Ib, 4.50 ± 0.81, one NMJ from each of 5 animals (N=5); *cac*^*sfGFP-N*^ Ib, 3.90 ± 1.07, N=8, p=0.67; wild-type Is, 6.28 ± 1.79, N=5; *cac*^*sfGFP-N*^ Is, 7.33 ± 2.20, N=6, p=0.72; Student’s t test). **(E)** Plot of the average amplitude of 1-sec, 20-Hz action potential train-mediated Ca^2+^ transients (wild-type Ib, 9.71 ± 1.53, one NMJ from each of 5 animals (N=5); *cac*^*sfGFP-N*^ Ib, 8.13 ± 2.21, N=8, p=0.57; wild-type Is, 12.26 ± 3.36, N=5; *cac*^*sfGFP-N*^ Is, 12.94 ± 3.45, N=6, p=0.89; Student’s t test). (**F-H**) Representative traces of EJPs and mEJPs recorded in 0.4 mM Ca^2+^ at wild-type (**F**), *cac*^*sfGFP-N*^ (**G**) and *cac*^*TagRFP-N*^ NMJs (**H**). **(I)** mEJP amplitude is unaffected in *cac*^*sfGFP-N*^ and *cac*^*TagRFP-N*^ (wild type, 0.91 ± 0.05, n=9 NMJs from 4 larvae; *cac*^*sfGFP-N*^, 0.99 ± 0.04, n=13 NMJs from 4 larvae, p=0.17, Student’s t test; *cac*^*TagRFP-N*^, 0.99 ± 0.04, n=12 NMJs from 4 larvae, p=0.20, Mann-Whitney U test). **(J)** EJP amplitude is unchanged between wild-type, *cac*^*sfGFP-N*^ and *cac*^*TagRFP-N*^ NMJs (wild type, 30.87 ± 1.41, n=9 NMJs from 4 larvae; *cac*^*sfGFP-N*^, 29.64 ± 0.52, n=13 NMJs from 4 larvae, p=0.84, Mann-Whitney U test; *cac*^*TagRFP-N*^, 26.70 ± 1.39, n=12 NMJs from 4 larvae, p=0.05, Student’s t test). **(K)** Quantal content is similar between wild-type, *cac*^*sfGFP-N*^ and *cac*^*TagRFP-N*^ NMJs (wild type, 34.8 ± 2.3, n=9 NMJs from 4 larvae; *cac*^*sfGFP-N*^, 30.6 ± 1.4, n=13 NMJs from 4 larvae, p=0.12, Student’s t test; *cac*^*TagRFP-N*^, 27.5 ± 1.7, n=12 NMJs from 4 larvae, p=0.03, Mann-Whitney U test).

### Endogenous Cac levels are heterogeneous and correlate with P_r_ at single AZs

*cac*^*sfGFP-N*^ allowed us to interrogate endogenous Ca^2+^channel levels and their relationship to single AZ function at the hundreds of individual synapses formed between a single motorneuron and postsynaptic muscle cell. Using Brp puncta to delineate single AZs, we found that Cac intensity per AZ varies considerably within a single motorneuron terminal, with most AZs exhibiting low-to-moderate levels and a smaller number expressing high Cac levels (Fig. 3A,B). This broad dynamic range in Ca^2+^ channel abundance could, in principle, underlie a similarly broad range of Ca2+ influx and P_r_ at individual AZs. Indeed, functional imaging studies at *Drosophila* NMJs have revealed significant heterogeneity in P_r_ between individual synapses of motorneuron-muscle pairs with the majority of AZs exhibiting low P_r_ and a small minority exhibiting high P_r_ – similar to the distribution we observe for Ca2+ channel levels (Guerrero et al., 2005; Peled and Isacoff, 2011; Melom et al., 2013; Peled et al., 2014; Akbergenova et al., 2017). As Ca^2+^ channel levels would be predicted to correlate with single-AZ P_r_ and a recent study observed a positive correlation between exogenous Cac levels and P_r_ (Holderith et al., 2012; Sheng et al., 2012; Akbergenova et al., 2017), we next examined the relationship between endogenous Cac levels and synapse-specific neurotransmitter release properties at motor AZs. We replaced sfGFP in the endogenous *cac* locus with sequence encoding the slow-bleaching, monomeric red-shifted fluorophore TagRFP and confirmed the tag did not disrupt synaptic transmission (See Fig. 2F-K). We then expressed postsynaptically targeted GCaMP5 (Akerboom et al., 2012) under the control of a muscle promoter (PS-GCaMP) in *cac*^*TagRFP-N*^ flies and monitored neurotransmission via Ca^2+^ influx through postsynaptic glutamate receptors during 1-Hz stimulation. Using nearest neighbor analysis, we assigned each Ca^2+^ influx event to an AZ defined by the center of mass of each Cac^TagRFP-N^-positive punctum and quantified the number of times a vesicle was released in response to 100 stimuli at individual AZs to calculate single-AZ P_r_ (Fig.s 3C,D; 3-1A-F). We observed significant heterogeneity in release probability, with a positively skewed distribution that matches observations from previous studies (Fig. 3-1G) (Peled and Isacoff, 2011; Melom et al., 2013; Peled et al., 2014). Specifically, we found that ∼90% of AZs had release probabilities between and.5, with a median P_r_ of.11, which corresponds remarkably well with physiological and prior optical imaging measurements (Lu et al., 2016; Newman et al., 2017).

When we compared P_r_ to Cac levels at individual AZs, we observed a highly positive correlation between Cac intensity and P_r_ as predicted (Fig. 3E-G). We considered the possibility that high-Cac, high-P_r_ AZs represented the combined molecular content and output of two adjacent AZs mistakenly identified as single AZs in our analysis. Our imaging parameters provided lateral resolution of ∼250 nm, so more closely spaced AZs would not be resolved. A recent study used super-resolution microscopy to investigate the spacing between AZs and determined that fewer than 2.5% of *Drosophila* motor AZs are separated by less than 280 nm (Akbergenova et al., 2017), indicating that the majority of high-P_r_/high-Cac AZs we observe are indeed single AZs. These data suggest the heterogeneity in Ca^2+^ channel levels is a key factor in establishing functional diversity between AZs of single neurons, and demonstrate that endogenously tagged Cac is a high-fidelity predictor of P_r._

**Figure 3.**
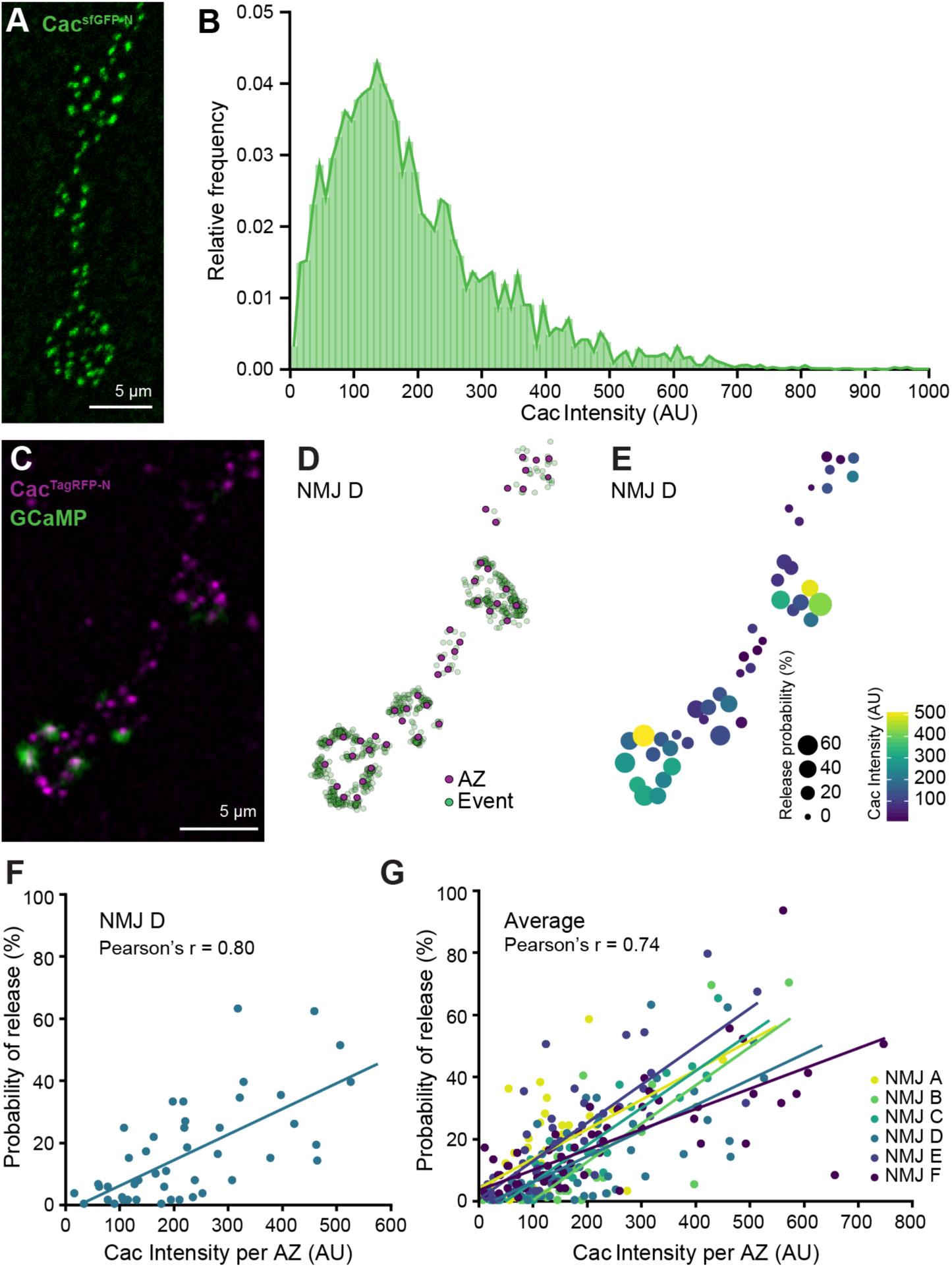
Cac^sfGFP-N^ is heterogeneously distributed and highly correlated with P_r_ at individual AZs. **(A)** Confocal Z-projection of a *cac*^*sfGFP-N*^ NMJ labeled with an antibody against GFP. **(B)** Broad distribution of Cac intensity across motorneuron AZs (n=3067 AZs [10 data points are outside the x-axis limits shown] from 16 NMJs of 4 animals). **(C)** Confocal Z-projection of Cac^TagRFP-N^ superimposed on a single frame of a PS-GCaMP movie monitoring neurotransmission in a live *cac*^*TagRFP-N*^; PS-GCaMP preparation. **(D)** Schematic of release events detected through postsynaptic GCaMP imaging (green dots) during 1-Hz stimulation mapped to individual AZs marked by Cac^TagRFP-N^ (magenta dots). **(E)** Heat map indicating the P_r_ and Cac intensity of each AZ by spot size and color, respectively. **(F)** Correlation between Cac and P_r_ at a single NMJ (NMJ D; shown in A-C) (Pearson correlation (r)=0.80, n=44 AZs). **(G)** Correlation between Cac and P_r_ at six NMJs in six larvae (average Pearson correlation (r)=0.74).

### Distinct regulation of Cac-sfGFP following the chronic expression of presynaptic homeostatic potentiation and depression

Various forms of adaptive plasticity operate to adjust presynaptic release properties (Regehr, 2012; Kittel and Heckmann, 2016; Jackman and Regehr, 2017; Monday and Castillo, 2017; Van Vactor and Sigrist, 2017). PHP is a conserved mechanism for maintaining synaptic strength within a stable range (Cull-Candy et al., 1980; Frank, 2014; Davis and Muller, 2015). At the *Drosophila* NMJ, genetic perturbation of postsynaptic glutamate receptors in *GluRIIA* loss-of-function mutants triggers a retrograde signal that instructs presynaptic neurons to chronically increase neurotransmitter release to precisely offset the reduced postsynaptic sensitivity to glutamate (Petersen et al., 1997). In contrast, presynaptic overexpression of the vesicular glutamate transporter vGlut increases the amount of glutamate contained in each synaptic vesicle. PHD restores synaptic strength through a decrease in the number of synaptic vesicles released per action potential (Daniels et al., 2004). Prior studies have revealed that increases in both Ca^2+^ influx and the readily releasable pool of synaptic vesicles (RRP) underlie PHP, whereas PHD is associated with a decrease in Ca^2+^ influx, but no change in the RRP (Weyhersmuller et al., 2011; Muller and Davis, 2012; Gavino et al., 2015; Kiragasi et al., 2017; Li et al., 2018a). Although the bi-directional modulation of Ca^2+^ influx contributes to the expression of both forms of homeostatic plasticity, the underlying mechanisms involved are not clear. In principle, bi-directional modulation of Ca^2+^ influx could be achieved through changes to the action potential waveform, Ca^2+^ buffering, Ca^2+^ channel gating properties, and/or Ca^2+^ channel abundance at AZs. A recent investigation ruled out modulation of the action potential waveform during both PHP and PHD (Gavino et al., 2015). However, it was recently reported that transgenically overexpressed Cac-GFP levels were decreased following overexpression of vGlut and increased in GluRIIA mutants (Gavino et al., 2015; Li et al., 2018a). Together, these studies suggest the simple model that bi-directional modulation of Ca^2+^ channel abundance drives opposing changes in Ca^2+^ influx during PHP and PHD. We tested this model using endogenously tagged Cac.

First, we demonstrated that both PHP and PHD are expressed normally in *cac*^*sfGFP-N*^ animals through intracellular recordings (Fig. 4A-F). Next, we investigated Cac^sfGFP-N^ levels in *GluRIIA* mutants to determine if endogenous Ca^2+^ channel levels are enhanced during chronic PHP expression. Consistent with the modulation of overexpressed Cac-GFP (Li et al., 2018a), we found that endogenous Ca^2+^ channel levels are indeed increased 36% in *GluRIIA* mutants (Fig. 4G-I). We also found that Brp levels were similarly increased (Fig. 4H, I) – a change that was previously reported and postulated to underlie the observed increase in RRP size (Weyhersmuller et al., 2011; Goel et al., 2017; Li et al., 2018a). Thus, a coordinated accumulation of both Brp and Cac are observed during chronic PHP expression, consistent with these AZ components contributing to the homeostatic increase in Ca^2+^ influx and the RRP.

**Figure 4.**
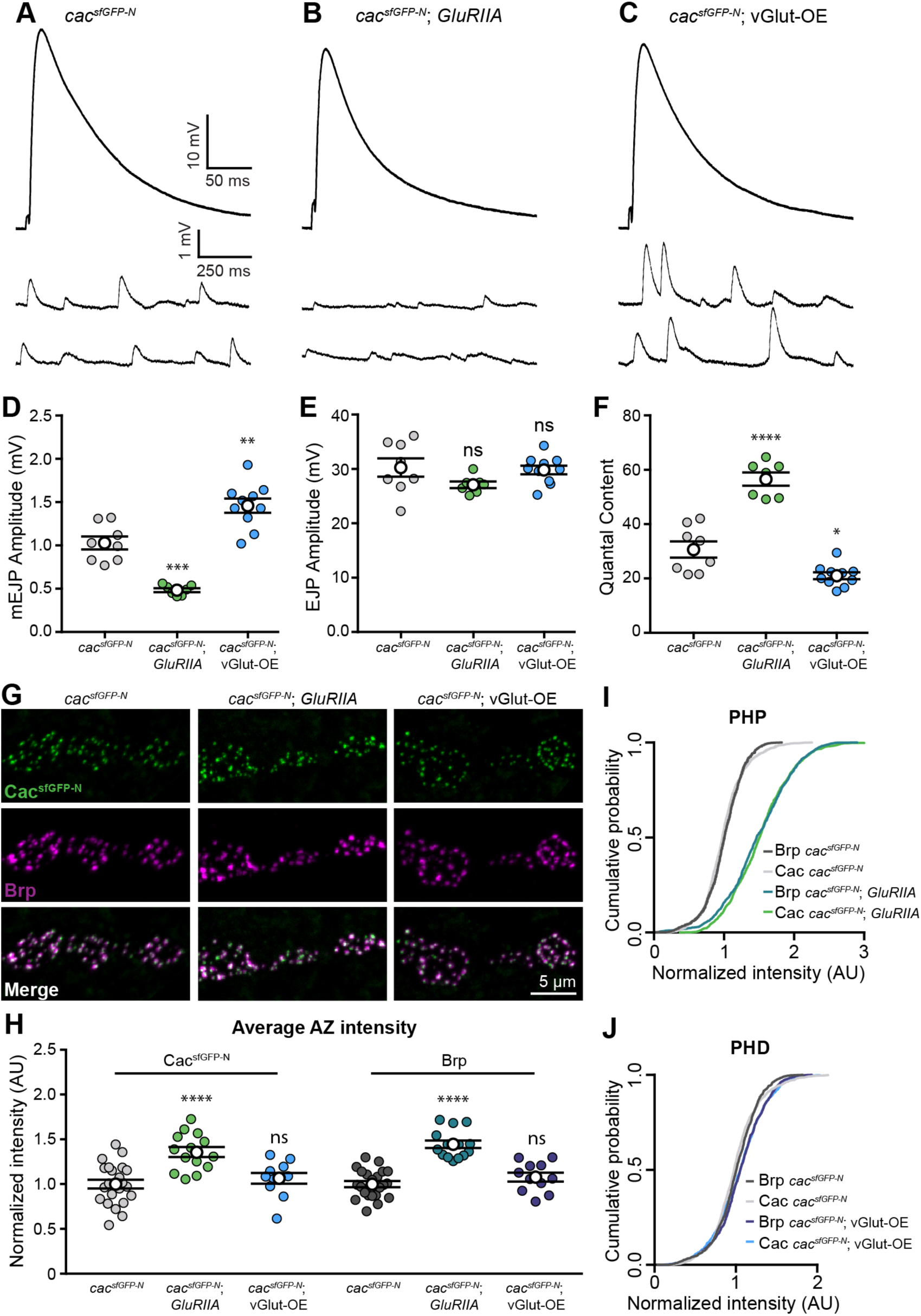
Endogenous Cac is differentially regulated in chronic PHP and PHD. (**A-F**) Chronic expression of PHP and PHD occurs normally in *cac*^*sfGFP-N*^ animals lacking GluRIIA or overexpressing vGlut, respectively. (**A-C**) Representative traces of EJPs and mEJPs recorded in 0.4 mM Ca^2+^ at *cac*^*sfGFP-N*^ (**A**), *cac*^*sfGFP-N*^; *GluRIIA* NMJs (**B**) and *cac*^*sfGFP-N*^; vGlut-OE (**C**). **(D)** mEJP amplitude is decreased in *cac*^*sfGFP-N*^; *GluRIIA* and increased in *cac*^*sfGFP-N*^; vGlut-OE as expected (*cac*^*sfGFP-N*^, 1.0 ± 0.08, n=8 NMJs from 4 larvae; *cac*^*sfGFP-N*^; *GluRIIA*, 0.48 ± 0.02, n=7 NMJs from 4 larvae, p=0.0001, t test with Welch’s correction; *cac*^*sfGFP-N*^; vGlut-OE, 1.5 ± 0.08, n=10 NMJs from 4 larvae, p=0.0017, Student’s t test). **(E)** EJP amplitude is unchanged between *cac*^*sfGFP-N*^, *cac*^*sfGFP-N*^; *GluRIIA* and *cac*^*sfGFP-N*^; vGlut-OE NMJs (*cac*^*sfGFP-N*^, 30.3 ± 1.70, n=8 NMJs from 4 larvae; *cac*^*sfGFP-N*^; *GluRIIA*, 27.1 ± 0.60, n=7 NMJs from 5 larvae, p=0.1115, t test with Welch’s correction; *cac*^*sfGFP-N*^; vGlut-OE, 29.8 ± 0.81, n=10 NMJs from 4 larvae, p=0.80, Student’s t test), as expected. **(F)** Quantal content is increased in *cac*^*sfGFP-N*^; *GluRIIA* and decreased in *cac*^*sfGFP-N*^; vGlut-OE as expected (*cac*^*sfGFP-N*^, 30.7 ± 3.0, n=8 NMJs from 4 larvae; *cac*^*sfGFP-N*^; *GluRIIA*, 56.6 ± 2.4, n=7 NMJs from 4 larvae, p<0.0001, Student’s t test; *cac*^*sfGFP-N*^; vGlut-OE, 21.0 ± 1.3, n=10 NMJs from 4 larvae, p=0.0145, t test with Welch’s correction). **(G)** Confocal Z-projections of Cac and Brp in *cac*^*sfGFP-N*^, in *cac*^*sfGFP-N*^; *GluRIIA* and *cac*^*sfGFP-N*^; vGlut-OE motorneuron boutons co-labeled with antibodies against GFP and Brp. **(H)** Normalized intensity of Cac and Brp puncta averaged for each NMJ in control, *GluRIIA* and vGlut-OE animals (Cac levels: *cac*^*sfGFP-N*^, 1.00 ± 0.05, n=22 NMJs from 9 larvae; *cac*^*sfGFP-N*^; *GluRIIA*, 1.36 ± 0.06, n=14 NMJs from 5 larvae, p<0.0001, Student’s t test; and *cac*^*sfGFP-N*^; vGlut-OE, 1.07 ± 0.06, n=11 NMJs from 5 larvae, p=0.43, Student’s t test. Brp levels: *cac*^*sfGFP-N*^, 1.00 ± 0.05, n=22 NMJs from 4 larvae; *cac*^*sfGFP-N*^; *GluRIIA*, 1.44 ± 0.04, n=14 NMJs from 5 larvae, p<0.0001, Student’s t test; *cac*^*sfGFP-N*^; vGlut-OE, 1.08 ± 0.06, n=11 NMJs from 5 larvae, p=0.19, Student’s t test). **(I)** Cumulative probability distributions of Cac and Brp puncta intensities demonstrate an increase in both Brp and Cac levels in *GluRIIA* animals (Cac levels: *cac*^*sfGFP-N*^, n=1,475 AZs; *cac*^*sfGFP-N*^; *GluRIIA*, n=1,475 - 3 data points are outside the x-axis limits shown. Brp levels: *cac*^*sfGFP-N*^, n=1,475 AZs; *cac*^*sfGFP-N*^; *GluRIIA*, n=1,475 AZs). **(J)** Cumulative probability distributions of Cac and Brp puncta intensities show no change in Brp or Cac levels in vGlut-OE animals (Cac levels: *cac*^*sfGFP-N*^, n=1,475 AZs; *cac*^*sfGFP-N*^; vGlut-OE, n=1,443 - 7 data points are outside the x-axis limits shown. Brp levels: *cac*^*sfGFP-N*^, n=1,475 AZs; *cac*^*sfGFP-N*^; vGlut-OE, n=1,443 AZs).

We next tested the model by investigating the modulation of endogenous Cac levels at NMJs overexpressing vGlut. In contrast to PHP, we did not observe a significant change in endogenous Ca^2+^ channel levels despite the robust expression of PHD (Fig. 4G-H,J). In agreement with Gavino et al. (2015), we also found no change in the abundance of the AZ cytomatrix protein Brp (Fig. 4G-H,J). To reconcile the difference between overexpressed and endogenous Cac, we repeated our analysis with Gal4-driven UAS-Cac-GFP. As previously reported, we observed robust PHD measured electrophysiologically together with a decrease in overexpressed Cac-GFP levels at AZs of type Is boutons, where Gavino et al. (2015) observed the greatest modulation (Fig. 4-1A-H). In contrast, we observed no change in endogenous Cac levels at AZs of type Is boutons overexpressing vGlut (Fig. 4-1H). This points to the use of the transgenic construct as the source of the discrepancy and highlights potential differences between the regulation of exogenous vs. endogenous ion channels. Together, these findings suggest that increased Cac abundance contributes to increased Ca2+ influx and P_r_ during chronic PHP adaptation, but that diminished Ca^2+^ influx during PHD is likely achieved through functional modulation of channel gating properties. More generally, these results indicate that AZ reorganization during PHP involves the coordinated recruitment of new vesicles to the RRP and an increase in Ca^2+^ channel levels, whereas PHD occurs without apparent morphological reorganization of the AZ, changes to RRP size, or modulation of Ca^2+^ channel levels – highlighting the diversity of responses employed by AZs to maintain stable synaptic communication.

### Ca^2+^ channel levels are rapidly increased at AZs during adaptation to acute postsynaptic receptor perturbation

At the *Drosophila* NMJ, PHP can be acutely induced through pharmacological inhibition of postsynaptic GluRIIA-containing glutamate receptors by the wasp venom philanthotoxin (PhTx). This triggers a potentiation of neurotransmitter release that precisely offsets the postsynaptic deficit within 10 minutes (Frank et al., 2006). There is evidence that the pathways that mediate chronic and acute PHP are partially overlapping, and both involve increased Ca^2+^ influx, RRP size, and Brp levels (Frank et al., 2006; Weyhersmuller et al., 2011; Muller and Davis, 2012; Muller et al., 2012; Goel et al., 2017; Kiragasi et al., 2017). However, the regulation of Cac levels during acute PHP expression has not been investigated.

To determine if endogenous Ca^2+^ channel levels are modulated over rapid timescales, we first assessed PhTx-induced PHP in *cac*^*sfGFP-N*^ larvae. As expected, we observe a robust homeostatic increase in neurotransmitter release (Fig. 5A-C). We then treated partially dissected *cac*^*sfGFP-N*^ larvae with nonsaturating concentrations of PhTx for 10 minutes and, immediately following PhTx treatment, fully dissected, fixed, and stained preparations for imaging of Cac and Brp. Strikingly, we observed a large increase in Cac levels (Fig. 5D-G). We also observed a significant increase in Brp levels across AZs (Fig. 5D-F,H) as previously reported (Weyhersmuller et al., 2011; Goel et al., 2017). Thus, Ca^2+^ channels accumulate at AZs within minutes during homeostatic potentiation of neurotransmitter release. Together with the prior findings that RRP levels increase on a similar timescale (Weyhersmuller et al., 2011), this indicates that AZs undergo a coordinated increase in size and molecular content during the acute expression of PHP, as they do over chronic timescales. Importantly, this finding does not rule out concurrent functional modulation of channels, but builds on prior findings to demonstrate that AZs are highly dynamic and capable of rapid growth that simultaneously increases not only the size of the RRP, but also Ca^2+^ channel levels.

**Figure 5.**
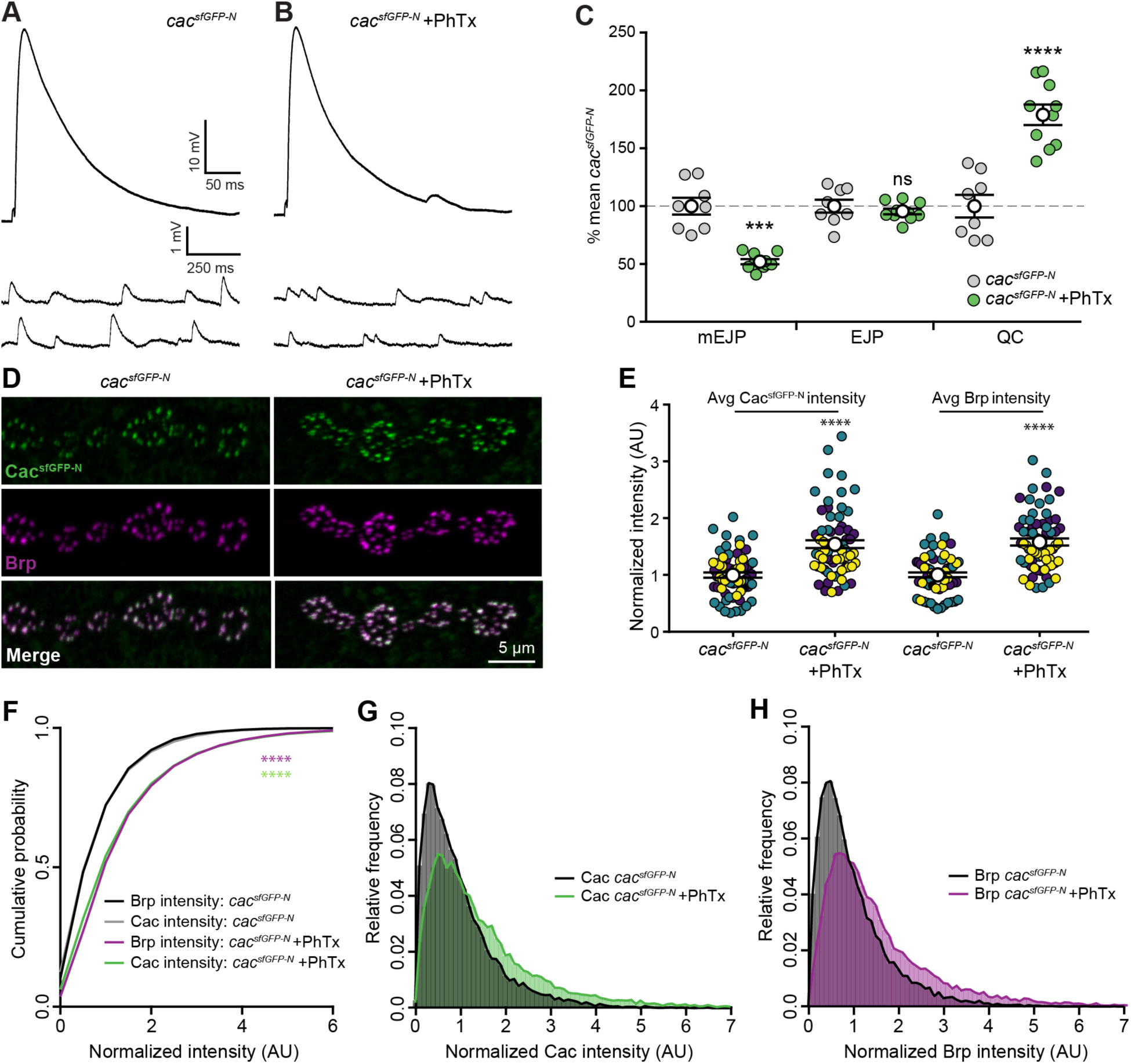
Cacophony levels are rapidly upregulated following acute homeostatic challenge. (**A,B**) Representative traces of EJPs and mEJPs recorded at *cac*^*sfGFP-N*^ NMJs in 0.4 mM Ca^2+^ with (**B**) or without (**A**) 10 min application of PhTx. **(C)** PhTx exposure significantly reduces mEJP amplitude at *cac*^*sfGFP-N*^ NMJs (*cac*^*sfGFP-N*^, 100 ± 7.3 %, n=8 NMJs from 4 larvae; *cac*^*sfGFP-N*^ + PhTx, 52 ± 2.2 %, n=10 NMJs from 4 larvae, p=0.0002, t test with Welch’s correction) as expected. In response to PhTx exposure, quantal content is significantly increased at *cac*^*sfGFP-N*^ NMJs (*cac*^*sfGFP-N*^, 100 ± 9.7 %, n=8 NMJs from 4 larvae; *cac*^*sfGFP-N*^ + PhTx, 179.1 ± 8.8 %, n=10 NMJs from 4 larvae, p<0.0001, Student’s t test), while EJP amplitude is maintained (*cac*^*sfGFP-N*^, 100 ± 5.6 %, n=8 NMJs from 4 larvae; *cac*^*sfGFP-N*^ + PhTx, 95.5 ± 2.5 %, n=10 NMJs from 4 larvae, p=0.4393, Student’s t test). **(D)** Confocal Z-projections of *cac*^*sfGFP-N*^ motorneuron boutons co-labeled with antibodies against GFP and Brp following vehicle (control) and PhTx treatment. **(E)** Normalized intensity of Cac and Brp per AZ averaged for each NMJ following 10-minute vehicle or PhTx treatment from three independent experiments (signified by yellow, green, and purple symbols) reveal a significant increase in both Brp and Cac levels immediately following PhTx treatment (Cac levels: control, 1.0 ± 0.05, n=65 NMJs from 18 larvae; PhTx, 1.54 ± 0.07, n=68 NMJs from 18 larvae, p<0.0001, Mann-Whitney U test. Brp levels: control, 1.0 ± 0.04, n=65 NMJs from 18 larvae; PhTx, 1.58 ± 0.06, n=68 NMJs from 18 larvae, p<0.0001, t test with Welch’s correction). **(F)** Cumulative probability distributions of Cac and Brp intensities in control and PhTx-treated animals (control, n=13,908 AZs; PhTx, n=15,925 AZs (80 data points are outside the x-axis limits shown), p<0.0001, Kolmogorov-Smirnov test). (**G,H**) Frequency distributions of Cac and Brp intensities at individual AZs of control and PhTx-treated animals reveal a rightward shift in intensities.

We observe significant differences in baseline Cac levels and P_r_ between individual AZs of single motorneurons. While the functional significance of this heterogeneity is unknown, it is clear that a small subset of AZs is responsible for the majority of neurotransmitter release under baseline conditions. PHP might similarly be achieved through the potentiation of a subset of AZs. Alternatively, potentiation may occur broadly across functionally heterogeneous AZs to scale release to an appropriate new level. We used live imaging of *cac*^*sfGFP-N*^ to address this question by quantifying Cac levels at the same AZs before and after PhTx-induced PHP. We observed increased Cac levels at identified AZs 10 minutes after PhTx application (Fig. 6A). Paired analysis of single AZs demonstrated that Ca^2+^ channel levels were significantly increased relative to control across AZs with heterogeneous baseline states, although levels change little or decrease at a subset of AZs (Fig. 6B-C). These findings are further supported by our fixed imaging results, where we observed a significant rightward shift in the entire distributions of Cac and Brp following PhTx treatment (See Fig. 5G,H). The smaller increase in Cac levels observed in our live imaging assays is likely due to the use of imaging conditions optimized for rapid acquisition and minimization of photobleaching. Given the heterogeneity in baseline Cac levels and P_r_, we next investigated how the degree of Ca^2+^ channel accumulation might be influenced by the baseline state of an AZ. Cac content could be increased by addition of the same amount of Cac to all AZs (additive model, green) or by an amount proportional to baseline levels (multiplicative model, orange). We found that the data is well explained by a multiplicative model, indicating a proportional increase in Ca^2+^ channel levels across heterogeneous AZs (Fig. 6D). We next examined the population of AZs where Cac content is unchanged or decreased, and found that they were more frequently observed among those AZs with lower baseline Cac levels (Fig. 6E). While imaging constraints make it harder to detect proportional changes in Cac levels at low-content AZs, we observe a significant effect consistent with the conclusion that baseline Cac content influences the likelihood of Ca^2+^ channel accumulation at a particular AZ. Notably, we did not observe significant formation of new AZs in our live preparations, a finding we confirmed in our fixed samples (Fig. 6F). This indicates that potentiation of release occurs largely through the strengthening of the majority of existing AZs. Together our findings provide insight into how functional heterogeneity is established at single AZs and further modulated during homeostatic plasticity.

**Figure 6.**
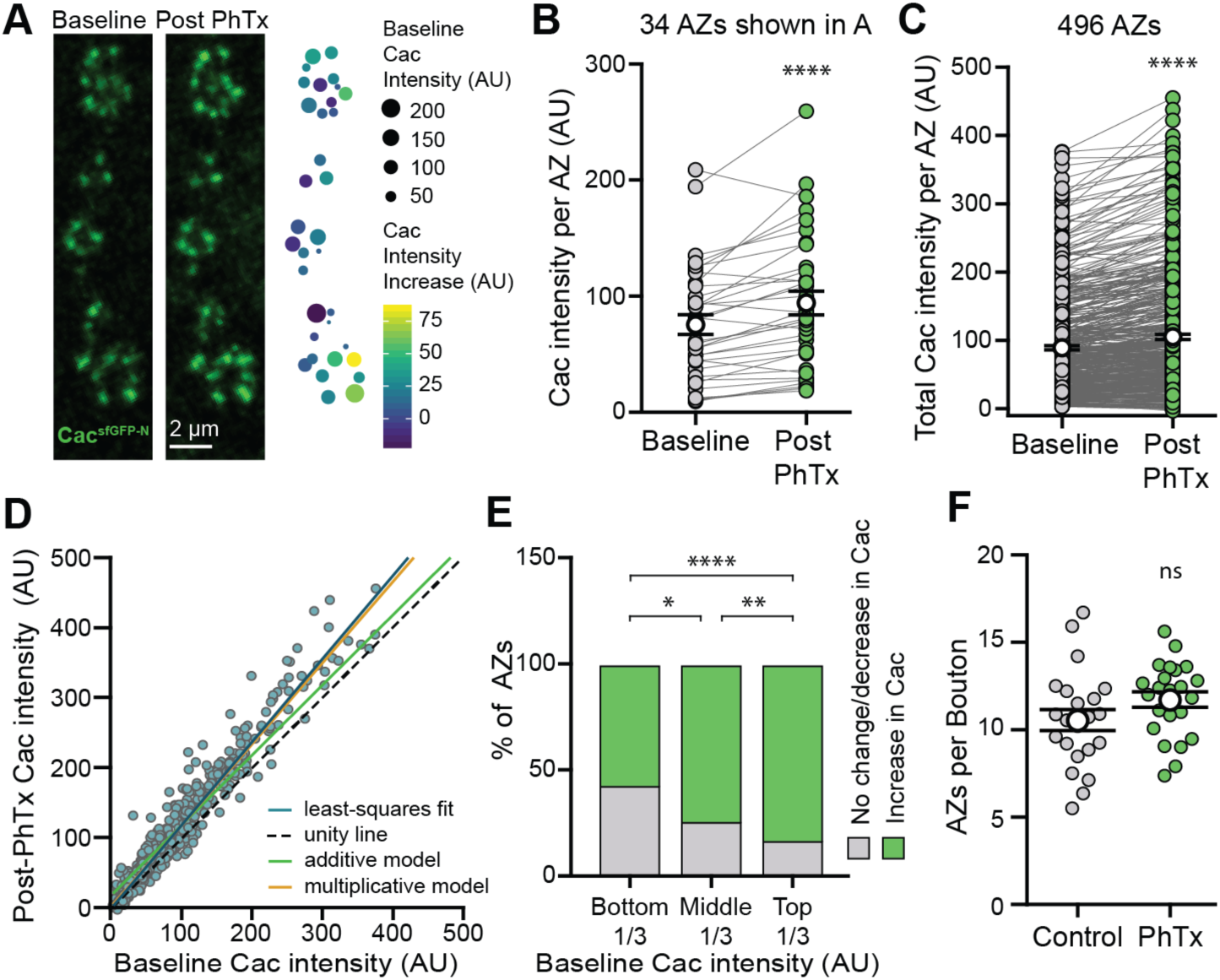
Live imaging of Cac^sfGFP^ reveals Ca^2+^ channel accumulation across heterogenous AZs during rapid homeostatic adaptation to PhTx application. **(A)** Confocal Z-projection of Cac at AZs of a motorneuron branch in a live *cac*^*sfGFP-N*^ preparation immediately before and 10 minutes after PhTx treatment. The heat map indicates PhTx-induced Cac accumulation (color) relative to baseline Cac levels (size) at each AZ. **(B)** Cac intensity at single AZs of the NMJ shown in (A) before and after PhTx reveals Cac accumulation at the majority of AZs (Baseline, 75.58 ± 8.55; Post PhTx minus vehicle, 94.25 ± 10.12, n=34 AZs from 1 NMJ branch, p<0.0001, Paired t test). **(C)** Cac intensity at single AZs across multiple animals also reveals Ca^2+^ channel accumulation at most AZs (Baseline, 89.56 ± 3.24; Post PhTx minus vehicle, 105.7 ± 3.96, n=496 AZs from 7 NMJs and 4 animals, p<0.0001, Wilcoxon signed rank test). **(D)** Cac intensity at individual AZs at baseline vs. post PhTx minus vehicle. The blue line is a least-squares fit to the data (slope =1.16, R^2^ = 0.94), while the dotted line would describe the data if there were no effect of PhTx (unity line, slope = 1), the green line would describe the data if there were an additive effect, and the orange line would describe the data if there were a multiplicative effect. **(E)** AZs that experience either no change or a decrease in Ca^2+^ channel content after PhTx occur more frequently in lower Cac-content AZs (bottom third, 48%; middle third, 28%, top third, 13%; bottom vs. middle, p=0.003, middle vs. top, p=0.02, bottom vs. top, p<0.0001, Fisher’s exact test). **(F)** AZ number is similar between control and PhTx-treated *cac*^*sfGFP-N*^ NMJs in fixed tissue preparations (control, 10.56 ± 0.60, n=22 NMJs from 6 animals; PhTx, 11.73 ± 0.45, n=23 NMJs from 6 animals, p=0.12, Student’s t test).

## Discussion

Diverse synaptic release properties enable complex communication and may broaden the capacity of circuits to communicate reliably and respond to changing inputs. We have investigated how the regulation of Ca^2+^ channel accumulation at AZs contributes to the establishment and modulation of AZ-specific release properties to maintain stable communication. Endogenous tagging of Cac allowed us to track Ca^2+^ channels live and in fixed tissue without the potential artifacts associated with transgene overexpression. This approach revealed differences in the regulation of endogenous and exogenous Ca^2+^ channels, underlining the value of developing and validating reagents for following endogenous proteins *in vivo*.

The abundance of endogenous Cac at individual AZs of single motorneurons is heterogeneous and correlates with single-AZ P_r_. This is consistent with previous studies in multiple systems linking endogenous Ca^2+^ channel levels at individual AZs to presynaptic release probability and efficacy, and a recent investigation of transgenically expressed Cac (Luo et al., 2011; Holderith et al., 2012; Sheng et al., 2012; Akbergenova et al., 2017). This strong correlation suggests Ca^2+^ channel levels might be regulated to tune P_r_ during plasticity, so we investigated the modulation of endogenous Cac levels in several *Drosophila* models of presynaptic homeostatic plasticity. Previous studies have suggested that the bi-directional regulation of Ca^2+^ influx at synapses contributes to the modulation of presynaptic release observed during both PHP and PHD (Frank et al., 2006; Muller and Davis, 2012; Muller et al., 2012; Gavino et al., 2015). A long-standing question is whether these changes are achieved through the regulation of channel levels, channel function, or through distinct mechanisms. Multiple mechanisms have been proposed to explain the increase in Ca^2+^ influx observed during the expression of PHP (Frank et al., 2006; Frank et al., 2009; Muller and Davis, 2012; Younger et al., 2013; Brusich et al., 2015; Muller et al., 2015; Kiragasi et al., 2017; Orr et al., 2017). For example, a presynaptic epithelial sodium channel (ENaC) and glutamate autoreceptor (DKaiR1D) have been implicated in promoting Ca^2+^ influx during PHP, leading to the model that modulation of presynaptic membrane potential might increase influx through Cac channels (Younger et al., 2013; Kiragasi et al., 2017; Orr et al., 2017). On the other hand, Frank and colleagues found that the guanine exchange factor Ephexin signals through the small GTPase Cdc42 to promote PHP in a Cac-dependent manner, raising the possibility that it does so through actin-dependent accumulation of channel levels (Frank et al., 2009). Further, multiple AZ cytomatrix proteins, including Fife, RIM, and RIM-binding protein, are necessary to express PHP and also regulate Ca^2+^ channel levels during development (Liu et al., 2011; Bruckner et al., 2012; Graf et al., 2012; Muller et al., 2012; Muller et al., 2015; Bruckner et al., 2017). However, whether Ca^2+^ channel abundance is modulated during PHP remained an open question.

Here, we demonstrate that Cac abundance is indeed enhanced during both the acute and chronic expression of PHP. This increase occurs in conjunction with the accumulation of Brp and enhancement of the RRP (Weyhersmuller et al., 2011; Goel et al., 2017; Li et al., 2018a), pointing to the coordinated remodeling of the entire neurotransmitter release apparatus during PHP on both timescales. Studies in mammals have found that AZ protein levels are dynamic and subject to homeostatic modification over chronic timescales (Matz et al., 2010; Lazarevic et al., 2011; Spangler et al., 2013; Glebov et al., 2017; Thalhammer et al., 2017), suggesting that structural reorganization of AZs is a conserved mechanism for modulating release. As ENaC and DKaiR1D-dependent functional modulation occur in tandem with the structural reorganization of AZs, it is interesting to consider why redundant mechanisms may have evolved. One remarkable feature of PHP is the incredible precision with which quantal content is tuned to offset disruptions to postsynaptic neurotransmitter receptor function. It is therefore tempting to hypothesize that PHP achieves this analog scaling of release probability by simultaneously deploying distinct mechanisms to calibrate the structure and function of AZs.

In contrast to the many mechanisms proposed for modulating Ca^2+^ influx during PHP, far less is known about how Ca^2+^ influx is regulated during PHD. One attractive idea was a reduction in AZ Ca^2+^ channel levels based on studies revealing reduced levels of transgenic UAS-Cac-GFP upon vGlut overexpression (Gavino et al., 2015). However, we found that although PHD is robustly expressed, endogenous Cac channels do not change in conjunction with vGlut overexpression. Because all Cac channels are tagged in *cac*^*sfGFP-N*^, this observation indicates that a reduction in Cac abundance at AZs is not necessary to achieve PHD. We determined that the source of the discrepancy is the use of the transgene to report overexpressed vs. endogenous Cac levels, demonstrating that exogenous and endogenous channels are regulated differently, at least during this form of PHD. This indicates that a mechanism other than modulation of Cac abundance drives PHD expression. Levels of Brp and RRP size are also unchanged during PHD (Gavino et al., 2015; Li et al., 2018a). Thus, the coordinated reorganization of the AZ appears to be specific to PHP. Interestingly, reversible downregulation of a subset of AZ proteins, but not Cac, was observed at *Drosophila* photoreceptor synapses following prolonged light exposure (Sugie et al., 2015). In the future, it will be of interest to determine if PHP and PHD share any mechanisms to control the bi-directional modulation of Ca^2+^ influx. PHD signaling operates independently of PHP, and was recently proposed to function as a homeostat responsive to excess glutamate, not synaptic strength, raising the possibility that mechanisms distinct from those that have been elucidated for PHP may regulate presynaptic inhibition during PHD (Li et al., 2018a).

Finally, live imaging of Cac^sfGFP-N^ during acute PHP enabled the investigation of how baseline heterogeneity in Cac levels and P_r_ intersects with the homeostatic reorganization of AZs. Monitoring endogenous Cac at the same AZs before and after PhTx treatment, we observed the accumulation of Ca^2+^ channels across AZs with diverse baseline properties. As with PHP expression over chronic timescales, our findings leave open the possibility of multiple mechanisms acting simultaneously, perhaps to ensure precise tuning, and do not rule out additional modulation of channel function or indirect regulation of Ca^2+^ influx. In fact, a prevailing model posits rapid events that acutely modulate P_r_ followed by consolidation of the response for long-term homeostasis (Frank et al., 2006; Frank et al., 2009). Coincident changes in Ca^2+^ channel function and levels coupled with long-term restructuring of AZs provides an attractive mechanism for this model. We also found that Cac accumulation is proportional across low-and high-P_r_ AZs. Therefore, baseline heterogeneity in Cac levels is maintained following the expression of PHP. At mammalian excitatory synapses, proportional scaling of postsynaptic glutamate receptor levels stabilizes activity while maintaining synaptic weights (Turrigiano, 2012). Our findings suggest an analogous phenomenon could be occurring presynaptically at the *Drosophila* NMJ. Notably, receptor scaling can occur globally or locally (Turrigiano et al., 1998; Yu and Goda, 2009). A recent study reported that PHP can be genetically induced and expressed within individual axon branches (Li et al., 2018b), demonstrating a similar degree of specificity in the expression of PHP at the *Drosophila* NMJ. A proportional increase in Cac levels could arise through homeostatic signaling from individual postsynaptic densities responding to similar decreases in quantal size – a strategy that would allow for both the remarkable synapse specificity and precision with which homeostatic modulation of neurotransmitter release operates.

## Supporting information

## Acknowledgements

We thank the Developmental Studies Hybridoma Bank, the Bloomington *Drosophila* Stock Center, and Aaron DiAntonio for antibodies and fly stocks. We are grateful to Fiona Ukken for generating postsynaptically targeted GCaMP5. We thank Maria Kamenetsky, UW-Madison CALS Statistical Consulting Lab, and Hong Zhan for assistance with data analysis and visualization. Elle Kielar-Grevstad, Director of the UW-Madison Biochemistry Optical Core, assisted with image analysis and Sean Carroll generously shared equipment. We thank Rick Ordway for his insightful comments. This work was supported by grants from the National Institute of Neurological Disorders and Stroke, National Institutes of Health to K.M.O.G (R01NS078179 and R21NS088830), D.D. (R01NS091546) and G.T.M. (R01NS061914), and a McKnight Technological Innovations in Neuroscience Award to K.M.O.G. S.J.G and K.M.O.G are currently affiliated with the Department of Neuroscience and Carney Brain Institute at Brown University. J.J.B is currently at the University of Oregon.

