## Supplementary material for "Endogenous tagging reveals differential regulation of Ca^2+^ channels at single AZs during presynaptic homeostatic potentiation and depression"

### Supplemental figures and legends

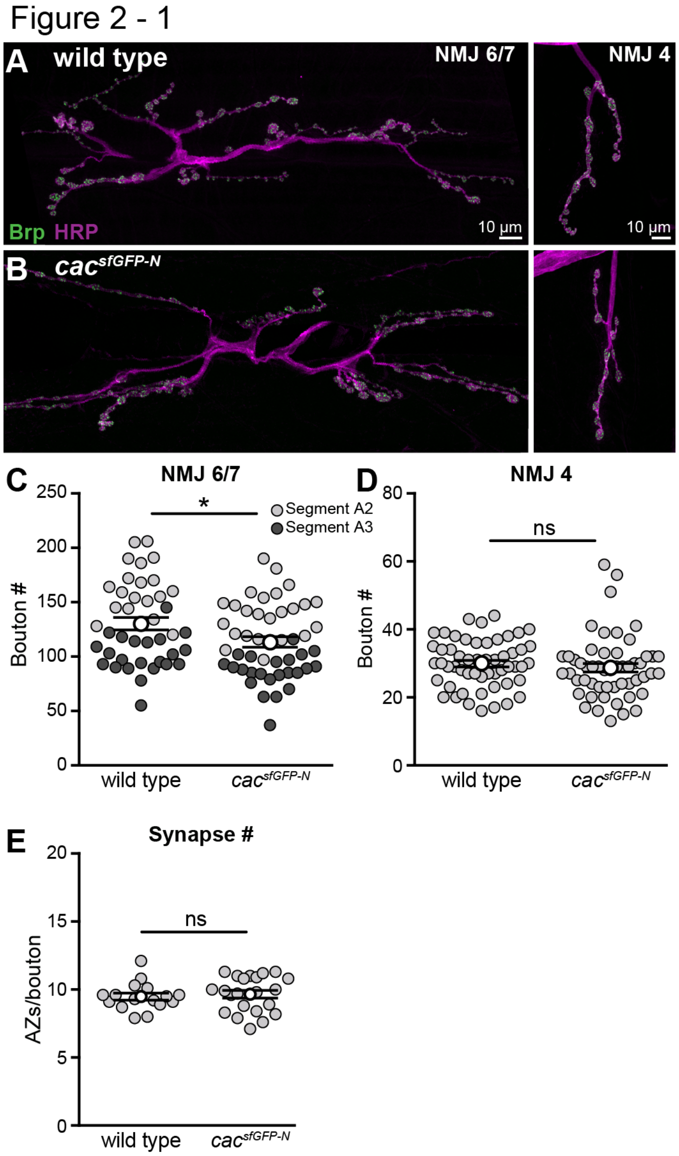


**Figure 2-1. Synaptic growth in *cac*$^{sfGFP-N}$.**

(**A,B**) Confocal Z-projections of NMJs 6/7 and 4 in wild-type and *cac*$^{sfGFP-N}$ larvae labeled with HRP to mark neuronal membranes and Brp to mark AZs.

(**C**) Quantification of bouton number at NMJ 6/7 reveals a small decrease in *cac*$^{sfGFP-N}$ (wild type, 130.2 ± 5.7, n=42 NMJs from 12 larvae; *cac*$^{sfGFP-N}$, 113.5 ± 4.8, n=48 NMJs from 13 larvae, p=0.026, Student's t test).

(**D**) Bouton numbers at NMJ 4 are similar in wild type and *cac*$^{sfGFP-N}$ (wild type, 29.9 ± 1.0, n=54 NMJs from 12 larvae; *cac*$^{sfGFP-N}$, 28.7 ± 1.3, n=53 NMJs from 13 larvae, p=0.15).

(**E**) AZ number is similar in wild type and *cac*$^{sfGFP-N}$ (wild type, 9.5 ± 0.26 AZs per bouton, n=16 NMJs from 9 larvae; *cac*$^{sfGFP-N}$, 9.7 ± 0.28 AZs per bouton, n=22 NMJs from 14 larvae, p=0.67, Student's t test).

Figure 3 - 1

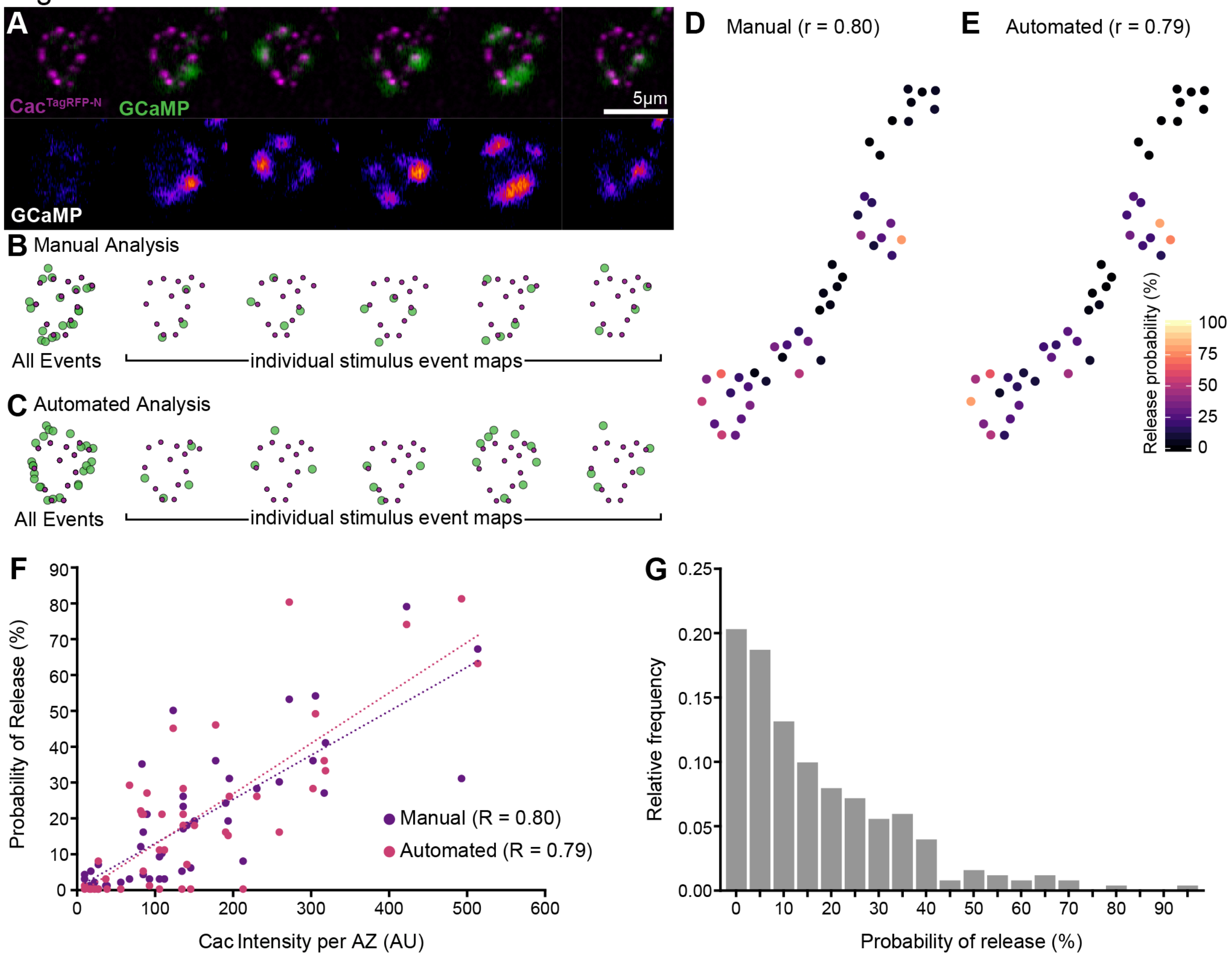


**Figure 3-1. Assignment of postsynaptic $Ca^{2+}$ transients to presynaptic AZs.**

(**A**) Top row: six frames of a GCaMP movie (one non-stimulus frame followed by five sequential stimulation frames) superimposed on a Z-projection of a live Z-stack of $Cac^{TagRFP\text{-}N}$ taken at the same terminal immediately before the movie. Bottom row: GCaMP channel from each frame shown separately using the "fire" look up table in FIJI.

(**B,C**) Illustration of manual and automated approaches for mapping $Ca^{2+}$ transients to AZs. XY coordinates of postsynaptic $Ca^{2+}$ transients on stimulation frames were gathered manually (**B**) or automatically (**C**) as displayed in the maps shown below each frame with $Cac^{TagRFP\text{-}N}$-marked AZs and $Ca^{2+}$ transient locations displayed as magenta and green circles, respectively (see 'Materials and methods' for details).

(**D,E**) Release probability maps resulting from manual (**D**) and automated (**E**) analysis.

(**F**) Correlation between Cac and $P_r$ using both methodologies (manual Pearson correlation (r)=0.80, automated Pearson correlation (r)=0.79).

(**G**) Distribution of $P_r$ across individual AZs of motor terminals synapsing on muscle 6 reveals a large number of low-$P_r$ AZs and small number of very high-$P_r$ AZs (n=251 AZs at 6 NMJs in 6 larvae).

Figure 4 - 1

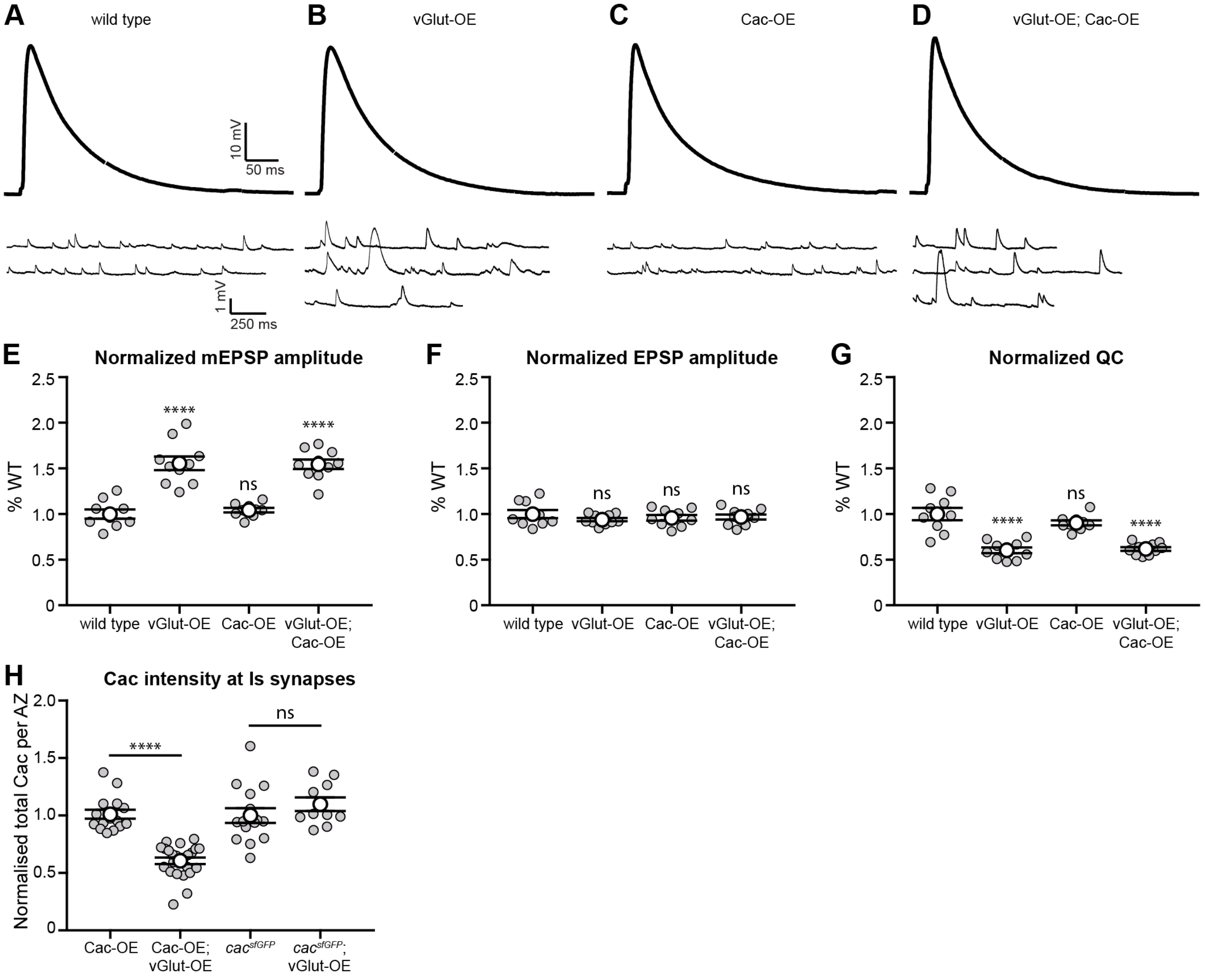


**Figure 4-1. Endogenous and transgenic Cac are differentially regulated during PHD.**

(**A-D**) Representative traces of EJPs and mEJPs recorded in 0.4 mM $Ca^{2+}$ at wild type (**A**), vGlut-OE (**B**), Cac-OE (**C**) and vGlut-OE; Cac-OE (**D**) NMJs.

(**E**) mEJP amplitude is increased in vGlut-OE and vGlut-OE; Cac-OE and unchanged in Cac-OE as expected (wild type, 1.0 ± 0.05, n=9 NMJs; vGlut-OE, 1.56 ± 0.07, n=10 NMJs, p<0.0001, one-way ANOVA; Cac-OE, 1.04 ± 0.02, n=9 NMJs, p=0.95, one-way ANOVA; vGlut-OE; Cac-OE, 1.55 ± 0.05, n=10 NMJs, p<0.0001, one-way ANOVA).

(**F**) EJP amplitude is unchanged between wild type, vGlut-OE, Cac-OE and vGlut-OE NMJs (wild type, 1.0 ± 0.05, n=9 NMJs; vGlut-OE, 0.94 ± 0.02, n=10 NMJs, p=0.53, one-way ANOVA; Cac-OE, 0.96 ± 0.03, n=9 NMJs, p=0.81, one-way ANOVA; vGlut-OE; Cac-OE, 0.97 ± 0.03, n=10 NMJs, p=0.88, one-way ANOVA).

(**G**) Quantal content is decreased in vGlut-OE and vGlut-OE; Cac-OE and unchanged in Cac-OE as expected (wild type, 1.0 ± 0.07, n=9 NMJs; vGlut-OE, 0.60 ± 0.03, n=10 NMJs, p<0.0001, one-way ANOVA; Cac-OE, 0.90 ± 0.03, n=9 NMJs, p=0.34, one-way ANOVA; vGlut-OE; Cac-OE, 0.62 ± 0.03, n=9 NMJs, p<0.0001, one-way ANOVA).

(**H**) Calcium channel abundance is decreased at Cac-OE*;* vGlut-OE Is NMJs but unchanged at *cac*$^{sfGFP\text{-}N}$*;* vGlut-OE Is NMJs (Cac-OE, 1.0 ± 0.04, n=15 NMJs; vGlut-OE; Cac-OE, 0.61 ± 0.03, n=24 NMJs, p<0.0001, Mann-Whitney U test; *cac*$^{sfGFP\text{-}N}$, 1.0 ± 0.06, n=15 NMJs; *cac*$^{sfGFP\text{-}N}$*;* vGlut-OE, 1.1 ± 0.06, n=10 NMJs, p=0.14, Mann-Whitney U test).
